# Food-web coupling by mobile consumers has individual to ecosystem level effects

**DOI:** 10.1101/2025.03.25.645357

**Authors:** Tianna Peller, Lara Chaouat, Henry Broadbent, Emanuele Giacomuzzo, Silvana Kaeser, George Mueller, Florian Altermatt

## Abstract

Mobile consumers serve as a ubiquitous link, coupling the dynamics and functions of different ecosystems, ranging from diel-vertical migration of zooplankton to seasonal movements of large ungulates. Despite mounting theoretical interest, experimental tests of the effects of consumer mobility in meta-ecosystems are virtually non-existent. Here, we used an experimental microcosm system to investigate how mobile consumers (*Daphnia magna*) mediate ecosystem biomass, community structure, and their own fitness by coupling spatially distinct aquatic ecosystems composed of two different protist communities. We found that consumer mobility significantly influenced the effect of consumption on ecosystem biomass and the growth and reproduction of the consumers, themselves. However, the direction and magnitude of these effects depended on community composition in the connected ecosystem. Further, we found consumer mobility consistently promoted the coexistence of a competing local species, regardless of community composition in the connected ecosystem. Our findings underscore the profound role of consumer mobility in shaping individual to ecosystem-level dynamics while emphasizing a strong mediating effect of community composition across the landscape.

## Introduction

Spatial structure plays a pivotal role in shaping ecological dynamics. The movement of organisms and resources across ecosystem boundaries establishes spatial connections that drive interactions and dependencies among ecosystems (Polis et al. 1997, Massol et al. 2011, Gounand et al. 2018a). These connections can influence species interactions, community structure, and ecosystem functioning on both local and landscape scales (McCann et al. 2005, Kremen et al. 2007, Bauer and Hoye 2014, Peller et al. 2024). While meta-ecology has traditionally focused on one type of organism movement connecting ecosystems—dispersal—recent calls have emphasized the urgent need for an improved integration and understanding of non-dispersal movements, highlighted both by theoretical and conceptual work in meta-ecosystem ecology (Gounand et al. 2018b, Guzman et al. 2019, Leroux and Schmitz 2025).

Mobile consumers represent a ubiquitous pathway by which ecosystems are connected in space (Lundberg and Moberg 2003). Emblematic examples include the diel-vertical migration of zooplankton, alternating between pelagic surface algal communities at night and deeper water heterotrophic plankton communities at day (Bandara et al. 2021), seasonal movement of large ungulates connecting different terrestrial ecosystems (Kauffman et al. 2021), to birds that connect terrestrial ecosystems to lakes and oceans (Green et al. 2023). As consumers tend to be more mobile than their prey, they commonly move across ecosystem boundaries to forage, coupling the dynamics of distinct ecosystems (Post et al. 2000, McCann et al. 2005, Peller et al. 2023). By coupling ecosystems in this manner, theory predicts that mobile consumers can stabilize food-webs, influence species coexistence, and drive spatially cascading effects, whereby distinct food-webs can influence one another via their effect on the shared consumer that connects them (de Roos et al. 1998, McCann et al. 2005; Rooney et al. 2008; Peller et al. 2022). While theory has greatly advanced our understanding of mobile consumer impacts, because it abstracts away ecological complexities, there remain substantial gaps in our knowledge of how these dynamics play out across real ecosystems: while large-scale movement and the crossing of ecosystems by mobile consumers is well documented (e.g., Bandara et al. 2021, Kauffman et al. 2021, Green et al. 2023), most of these examples are observational and do not allow replicated experimental testing (but see Little et al. 2019). Such experimental research can provide a crucial complement to develop our understanding by testing hypotheses and generating new predictions through controlled yet ecologically relevant experiments (Lawler 1998, Srivastava et al. 2004, Altermatt et al. 2015), as has been done in local ecosystem contexts (e.g., Leibold 1989, Fussman et al. 2000, Petchey 2001). However, despite mounting interest in the ecological role of mobile consumers, experimental tests of consumer-mediated effects across coupled ecosystems remain absent.

An empirical feature that theory has largely overlooked is the role of community diversity in shaping the impacts of consumer mobility. Over the past few decades, observational evidence has revealed a great deal about how mobile consumers move across landscapes and interact with diverse ecological communities (Meyer et al. 1983, Abbas et al. 2012, Donadi et al. 2017, Peller et al. 2021). For example, birds forage in both marine and terrestrial ecosystems, which typically have limited overlap in species composition, while bears regularly move between rivers and forests to forage (Deacy et al. 2017), and zooplankton interact with the different communities of the benthic and pelagic realms (Bandara et al. 2021). Local scale experiments have shown the dynamics and impacts of consumers within an ecosystem are significantly shaped by the characteristics of the prey community in the ecosystem, including prey availability, diversity, and composition (Petchey 2001). Extending this concept to mobile consumers suggests that the composition of prey in one ecosystem may indirectly shape the dynamics of other ecosystems through the shared consumer that connects them.

Additionally, field evidence suggests that where mobile organisms feed can directly influence their fitness, further underscoring the importance of community diversity in shaping consumer effects. If the fitness of mobile consumers is tied to the specific prey communities they exploit, then differences in prey availability across ecosystems may have cascading effects on both the consumers and the ecosystems they connect. However, theory has yet to fully integrate these insights, leaving a significant gap in our understanding of how mobile consumers mediate cross-ecosystem dynamics. As global change continues to alter both species movement patterns and community structures, addressing this gap is increasingly urgent (Rosenberg et al. 2019, Williams et al. 2021).

Here, we used an experimental microcosm system to test the impacts of mobile consumers and the role of community diversity, using mobile *Daphnia* grazers feeding on differently composed plankton communities including ciliates and rotifers. While not aimed to directly mimic the diel-vertical movement of *Daphnia* and the different plankton communities they graze on, the system contains organisms that are realistically involved in such large-scale cross-ecosystem movement scenarios in natural systems. We built two-patch meta-ecosystems consisting of aquatic microbial communities, with all meta-ecosystems exhibiting the same community composition in ecosystem 1, but differences in community composition in ecosystem 2 (Fig. 1). We carried out controlled movements of mobile consumers (*Daphnia magna*) between ecosystem 1 and 2 for a total of three complete return movements. Our findings show that consumer mobility significantly reduced the negative impact of consumers on ecosystem biomass, but that the composition of the connected community could mediate the magnitude of this effect. We found consumer mobility could significantly influence consumer growth and reproduction, but similarly that community composition had a mediating effect on the direction and magnitude. In addition, we found consumer mobility consistently promoted the coexistence of a competing local species. Ultimately, our experiment demonstrates the effects of consumer mobility on ecosystems, and on the consumers themselves, can be strongly shaped by the composition of the ecosystems they connect.

**Figure 1.**
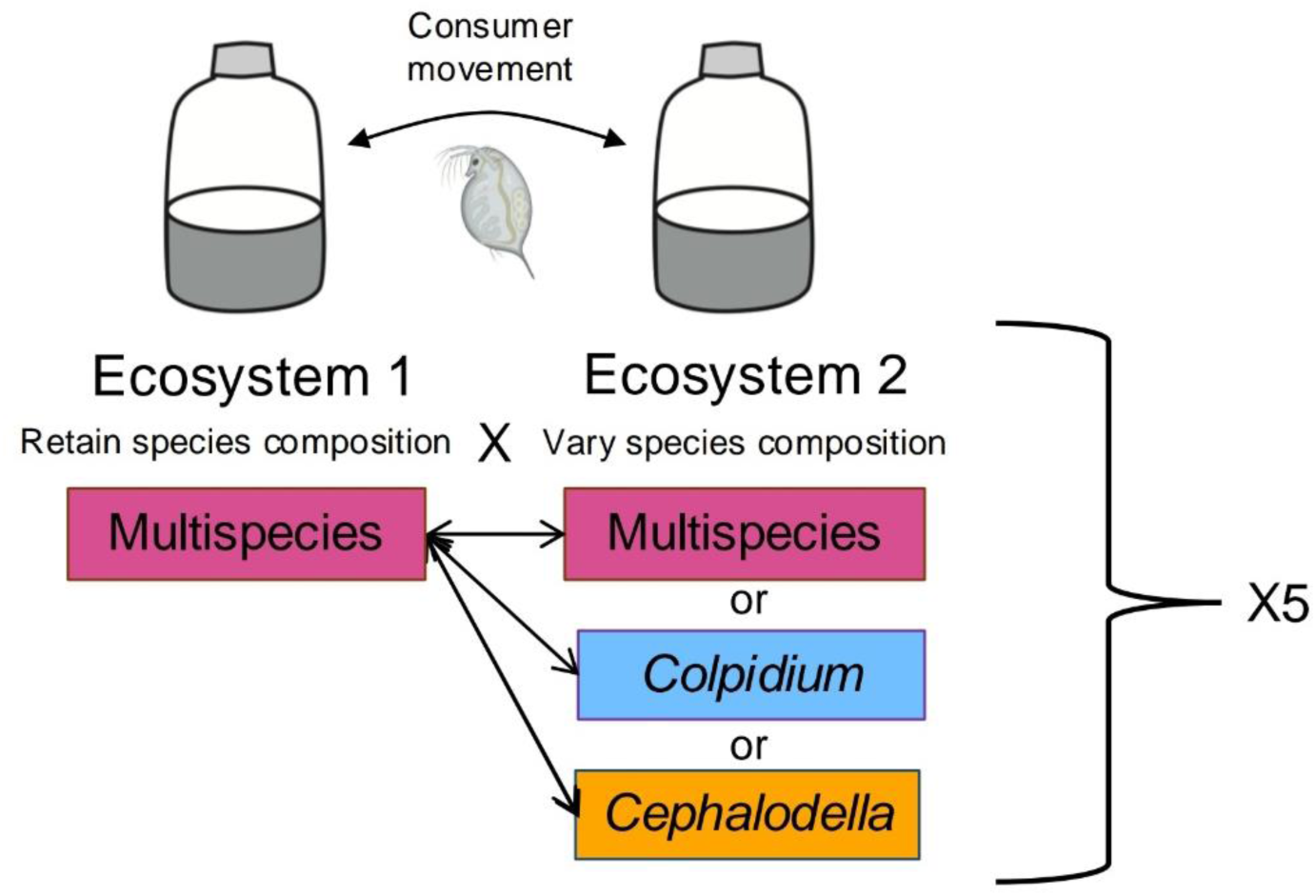
The experimental design consists of two-patch meta-ecosystems connected via the movement of consumers (*Daphnia magna*). Ecosystem 1 exhibits the same 10-species mixed composition across treatments (Multispecies; pink). In ecosystem 2, we varied the species composition to include either 1) the same 10-species mixed composition as ecosystem 1 (Multispecies; pink), 2) a *Colpidium striatum* monoculture (*Colpidium*; blue), or 3) a *Cephalodella* sp. monoculture (*Cephalodella*; yellow). This combination of ecosystem 1 and the three variations of ecosystem 2 gave three different meta-ecosystems, each of which was replicated five times. Isolated controls with and without consumers (not shown) exhibited the same 10-species mixed composition and were similarly replicated five-fold.

## Methods

### Experimental design

We studied the influence of mobile consumers and the potential mediating role of local community composition using a protist microcosm experiment (Altermatt et al. 2015). Each individual microcosm represented a distinct ecosystem, with either a mixed community, or a monoculture of a single species present within the mixed community. The experiment included three meta-ecosystem variations, each consisting of two microcosms connected via the controlled movement of mobile consumers (Fig. 1): 1) multispecies meta-ecosystems, consisting of two microcosms with the mixed multispecies community; 2) multispecies-*Colpidium* meta-ecosystems, consisting of one microcosm with the mixed multispecies community and one microcosm with a *Colpidium* sp. monoculture; and 3) *multispecies-Cephalodella* meta-ecosystems, consisting of one microcosm with the mixed multispecies community and one microcosm with a *Cephalodella* sp. monoculture. In addition, the experiment consisted of two groups of isolated control ecosystems: 1) multispecies isolated ecosystems without consumers, which consisted of a single microcosm with the multispecies community and the mobile consumer species absent; and 2) multispecies isolated ecosystems with consumers, which consisted of a single microcosm with the multispecies community and the mobile consumer species present, but restricted to the single microcosm.

We used *Daphnia magna* as mobile consumers in our experiment. There is abundant evidence that *Daphnia* prey on protists and rotifers. For instance, field studies have demonstrated that *Daphnia* can suppress ciliate populations, even at densities as low as <1 individual per liter (Wickham and Gilbert, 1991, Zöllner et al. 2003), indicating that some natural ciliate populations are highly susceptible to *Daphnia* predation (Pace and Funke 1991; Wickham and Gilbert 1991). Notably, we selected the monoculture species present in microcosms (ecosystem 2; the ciliate *Colpidium* sp. and the rotifer *Cephalodella* sp.) based on the size of the species, as *Daphnia* are known to be more efficient at consuming small prey (McMahon and Rigler 1965; Tezuka, 1974; Porter et al., 1979; DeBiase et al., 1990; Jack and Gilbert, 1993). The *Colpidium* monoculture, thus, represented a small (∼45 μm long; Giometto et al. 2013), easier to consume prey, while the *Cephalodella* monoculture represented a larger (∼150μm long), more difficult to consume prey. For all meta-ecosystems, we performed controlled movements of *Daphnia* between the two local ecosystems every 2.5 days for a total of three complete return movements. For isolated ecosystems with consumers, *Daphnia* remained in the isolated ecosystem for the duration of the experiment. The experiment was replicated five-fold.

### Experimental setup

The mixed communities consisted of seven heterotrophic ciliates: (*Blepharisma* sp., *Colpidium* sp., *Loxocephalus* sp., *Paramecium aurelia*, *Paramecium caudatum*, *Spirostomum teres*, and *Tetrahymena* cf. *pyriformis*), two mixotrophic ciliates able to photosynthesise (*Euglena gracilis* and *Euplotes aediculatus*), and one rotifer (*Cephalodella* sp.). We subsequently refer to all species in the mixed community as ‘protists’. We cultured the protists in pre-autoclaved bottles with standard protist medium (0.46 g of Protozoa Pellet by Carolina per L of water) and a bacterial culture (*Serratia fonticola*, *Bacillus subtilis*, and *Brevibacillus brevis*) serving as food for protists and constituting 5% of the total culture volume. Further details on the protist culture and experimental procedures can be found in Altermatt et al. (2015).

At the beginning of the experiment (day zero), we prepared a master mix of the mixed community by combining all 10 species at 1/10 of their carrying capacity, with the mixture volumetrically supplemented with 15% standard protist medium. We similarly prepared a master monoculture for *Colpidium* and *Cephalodella*, from which the monoculture ecosystems were filled. The experiment was conducted in 100 mL Schott bottles, with each bottle representing an ecosystem. We pipetted 50 mL of the master mix to all multispecies ecosystems. Likewise, we pipetted 50 mL of the *Colpidium* and *Cephalodella* monoculture to their single-species ecosystems, respectively.

On day zero of the experiment, two *Daphnia* of similar size were added to ecosystem 2 in the meta-ecosystems and to the isolated control ecosystems with consumers, where they remained until the first consumer transfer day. *Daphnia* were added using glass pipettes. A photograph of each *Daphnia* was taken before they were added to their respective ecosystem to enable size measurements (see following section). The replicates were randomised in position and kept in an incubator at 20 °C with constant lighting for the remainder of the experiment.

### Consumer movement, size measurement, and reproductive output

Every 2.5 days, both *Daphnia* were transferred between Schott bottles, which represented the movement of mobile consumers between ecosystems. Using glass pipettes, both *Daphnia* were pipetted with into a single petri dish. While in the petri dish, we took photographs of each *Daphnia*, individually, using a microscope with a 1.6x magnification and a Canon D5126321 camera with a 2.5x magnification. To avoid protists being transferred between ecosystems during this process, *Daphnia* were subsequently washed using lake water before being returned to the opposite Schott bottle in the meta-ecosystem pair. *Daphnia* in the isolated systems were similarly removed from the Schott bottle, photographed and washed; however, *Daphnia* from the isolated systems were subsequently returned to the same Schott bottle. For isolated systems without *Daphnia*, we introduced the same handling treatment to the system by pipetting out and subsequently returning a small volume of lake water to the system.

The size of each *Daphnia* at each transfer was determined using the photographs taken with the camera and analyzed using the software ImageJ (Schneider et al. 2012). Specifically, we measured the body length of all *Daphnia*, defined as the diagonal measurement from the top of the eye to the apical base of the spine (Duckworth et al. 2019). We counted *Daphnia* offspring visually during each transfer event and on the final day of the experiment. *Daphnia* offspring were removed from each ecosystem and discarded to maintain consistency in the experiment and avoid confounding effects. By removing offspring, we ensured that the population size of the *Daphnia* remained stable across the experimental timeline, allowing us to focus on the effects of adult consumer mobility.

### Sampling

To determine species identities, species biomass, and traits of protists in each ecosystem, we took videos of each ecosystem every 2.5 days, during each predator transfer event, as well as on the first and final day of the experiment. Specifically, we took a 5 s-video of 0.2 mL samples at 1.6x magnification, using a Hamamatsu Orca Flash 4.0 (Herrsching am Ammersee, Germany) camera. Prior to the start of the experiment, we recorded videos of all protist monocultures to create a training dataset for species identification based on their traits. For each monoculture, we captured enough footage to include at least 100 individuals per species.

### Quantifying biomass and species dominance

We used the R-package BEMOVI to identify and characterise protist species in each ecosystem (Altermatt et al. 2015; Pennekamp et al. 2015). From each video, we extracted moving particles’ traits (e.g., speed, shape, size), which we used to filter out particles that were not protists and obtain an average abundance of protist individuals per volume. We calculated the total area of protists (as area per volume medium), which we used as a proxy of biomass (and hereafter refer to as biomass) following previous studies (e.g., Pennekamp et al. 2018, Giacomuzzo et al. 2024). We identified protist species with a support vector machine model (Cortes et al. 1995; r-package “e1071”: Dimitriadou et al. 2006), using traits extracted from species monocultures as predictor variables. Using the species identifications obtained, we calculated species dominance based on the total biomass of protists.

### Statistical analysis

We determined the effect of consumer mobility and community composition in ecosystem 2 on total biomass in ecosystem 1 using log response ratios (LRR). Specifically, we used LRR of protist biomass in ecosystem 1 at each time point of the experiment, for the three meta-ecosystems and the isolated ecosystem with *Daphnia* consumers, with 95% confidence intervals. We tested the responses relative to the isolated control without *Daphnia* consumers, such that confidence intervals not overlapping with zero reveal significant effects of consumption. Confidence intervals for meta-ecosystems that do not overlap with confidence intervals from the isolated control with *Daphnia* consumers indicate a significant effect of consumer mobility, while non-overlapping confidence intervals of different meta-ecosystem treatments indicate a significant effect of initial community composition in ecosystem 2. Positive (negative) LRR values indicate *Daphnia* consumption had a positive (negative effect) on biomass in ecosystem 1. We studied the potential interaction with time by running generalized linear models (GLM) with Gaussian distributions as link functions, on LRRs with treatment and time as explanatory variables. We assessed significance and effect size of each factor, and their interaction, using a type II analysis of deviance. We performed post hoc analyses to evaluate significance of pairwise differences between treatments (with Tukey adjusted p-values).

To assess differences in species dominance across treatments at the end of the experiment in ecosystem 1, we calculated Bray-Curtis dissimilarity indices and performed a permutational multivariate analysis of variance (PERMANOVA). Specifically, we computed a Bray-Curtis dissimilarity matrix to quantify differences in community composition in ecosystem 1 across treatments, with regard to dominance based on species biomass. To test for significant differences between meta-ecosystem types, we performed PERMANOVA, with meta-ecosystem type as the primary predictor. Statistical significance was determined based on permutation tests, with results reported as F-values and associated p-values.

We determined the effect of consumer mobility and community composition in ecosystem 2 on the size of consumers using mixed effects models. Specifically, we tested the interactive influence of consumer treatment and continuous time on consumer size, with replicate as a random effect. As each replicate had two consumers, the response variable was the mean size per replicate. Male consumers were excluded from the analysis, as our focus was on reproductive potential and the potential for intergenerational effects. We performed post hoc analyses to evaluate significance of pairwise differences between treatments (with Tukey adjusted p-values).

We investigated the effect of consumer mobility and community composition in ecosystem 2 on the reproductive output of consumers using an analysis of variance (ANOVA). A one-way ANOVA was conducted to assess whether the number of offspring per consumer differed significantly across treatments. For replicates with two females, the total offspring count was divided by two to give a mean number of offspring per female. In replicates with one male and one female, no division was performed, as only the female’s offspring production was included. This approach ensured that all replicates were standardized to account for differences in sex composition. We performed post hoc pairwise comparisons using Tukey’s Honest Significant Difference test to identify specific treatment differences.

All analyses were conducted with R version 3.1.2, with packages ‘car’ (Fox and Weisberg 2011) for type II analysis of deviance, ‘lme4’ (Bates et al. 2015) for mixed effect models, ‘emmeans’ (Lenth 2022) for post hoc comparisons, and ‘vegan’ (Oksanen et al. 2024) for computing Bray-Curtis dissimilarity matrices. Analyses across time did not include the initial two time points before the consumer movement began (time = 0, 2), as they are not related to the understanding of the effects of consumer movement.

## Results

### Consumer mobility and composition of the connected ecosystem affect ecosystem biomass

We found that both the mobility of the consumer and the community composition of the connected ecosystem (ecosystem 2) significantly influenced total biomass in ecosystem 1 (Fig. 2, Chisq = 20.57, df = 3, p < 0.001, Table S1), with a significant interaction with time (Chisq = 10.93, df = 3, p = 0.012, Table S1). Relative to the immobile consumer treatment, where consumers were restricted to a single ecosystem, connecting the consumers to a second ecosystem with the same community composition (multispecies meta-ecosystem) significantly reduced the negative effect of consumption on biomass across the experiment (Fig. 2; pink versus grey points; t = −3.112, df = 12, p = 0.039). However, we observed that this influence of consumer mobility on ecosystem biomass could be mediated by the community composition in the connected ecosystem. Being connected to a *Cephalodella* monoculture led to strong negative impacts on biomass in ecosystem 1 directly following consumer visits, such that it was not significantly different from the immobile consumer treatment across the experiment (Fig. 2; yellow points versus grey points; t = −0.894, df = 12, p = 0.81). Whereas, being connected to a *Colpidium* monoculture led to a similar impact on biomass in ecosystem 1 relative to the multispecies meta-ecosystem (Fig. 2; blue points versus pink points; t = 0.196, df = 12, p = 0.83), although tended to mildly reduce the negative impact. Consequently, biomass in ecosystem 1 was significantly higher when consumers were connected to a *Colpidium* monoculture compared to a *Cephalodella* monoculture (Fig. 2; blue versus yellow points; t = 3.063, df = 12, p = 0.043). In Figure S1, we show that biomass in ecosystem 2 did not differ significantly among meta-ecosystem types, suggesting these results were not driven by quantitative differences in resource availability.

**Figure 2.**
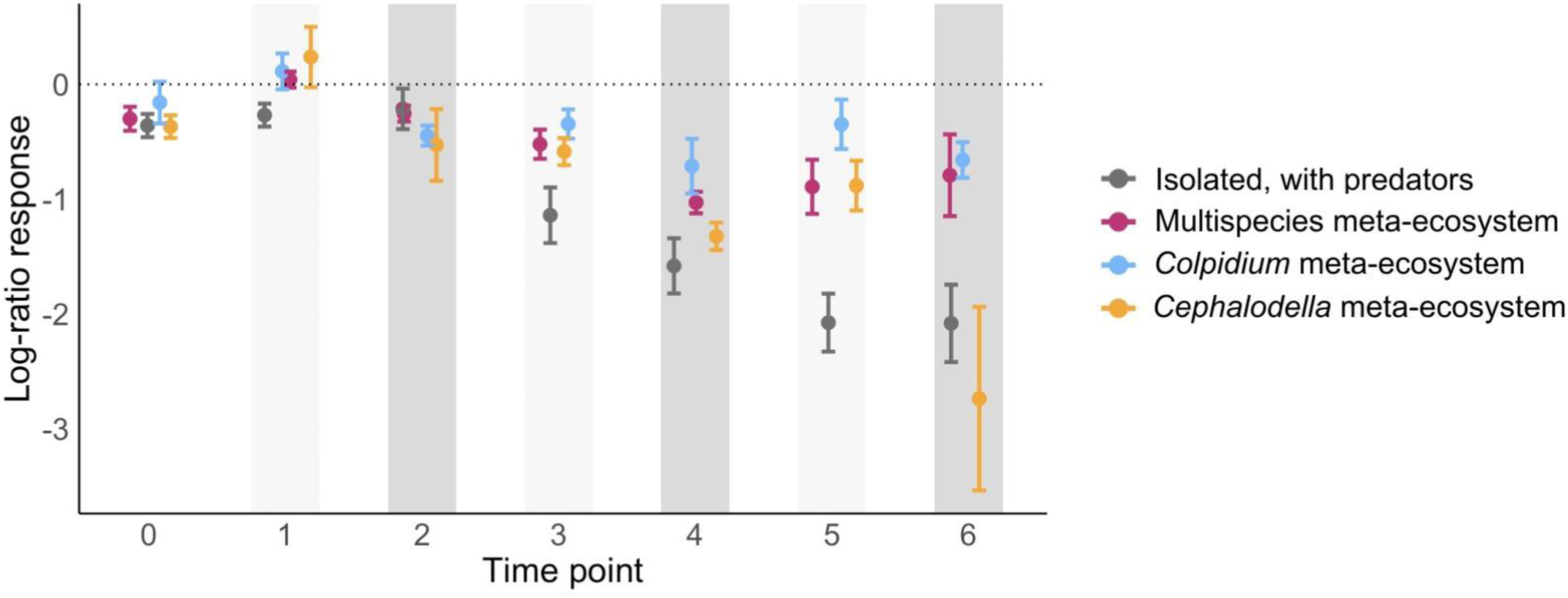
Effects of *Daphnia* predation on the total biomass of the multispecies community in ecosystem 1 over time. Effects are expressed as the log response-ratio (LRR) of total biomass in ecosystem 1 in treatments with *Daphnia* consumers compared to the isolated control without *Daphnia*. Colors of points indicate the species composition in the connected ecosystem (ecosystem 2); grey points show the isolated ecosystems with *Daphnia* constantly present. Bars indicate the 95% confidence intervals. Confidence intervals not crossing the zero dotted line differ significantly from the control, with no *Daphnia* predators. Non-overlapping confidence intervals between treatments reveal that the effect significantly differs between the treatments. Time points with light grey shading show the biomass in ecosystem 1 directly before *Daphnia* consumers were transferred to ecosystem 1. Whereas time points with dark grey shading show the biomass in ecosystem 1 directly after *Daphnia* consumers were removed from ecosystem 1 and transferred to ecosystem 2. Time point zero is not shaded as it represents the start of the experiment before any *Daphnia* had been added.

### Consumer mobility affects species dominance rankings

We found that species dominance in ecosystem 1 at the end of the experiment depended on the consumer treatment (Fig. 3; *F*_4, 20_ = 1.98, p = 0.034). In particular, we observed significant effects of consumer mobility on the dominance of *Cephalodella* sp. (Cep) (Fig. 3, light green). In the isolated ecosystem without consumers and in the mobile consumer treatments, the same two species consistently dominated the system: *Cephalodella* sp. (Cep) and *E. aediculatus* (Eup). However, in the isolated ecosystem with immobile *Daphnia* consumers, *Cephalodella* sp. was excluded from the community, reaching zero by the end of the experiment. Thus, a different dominance ranking was observed: *E. aediculatus* was the most dominant species, and *Spirostomum teres* was the second most dominant species at the end of the experiment.

**Figure 3.**
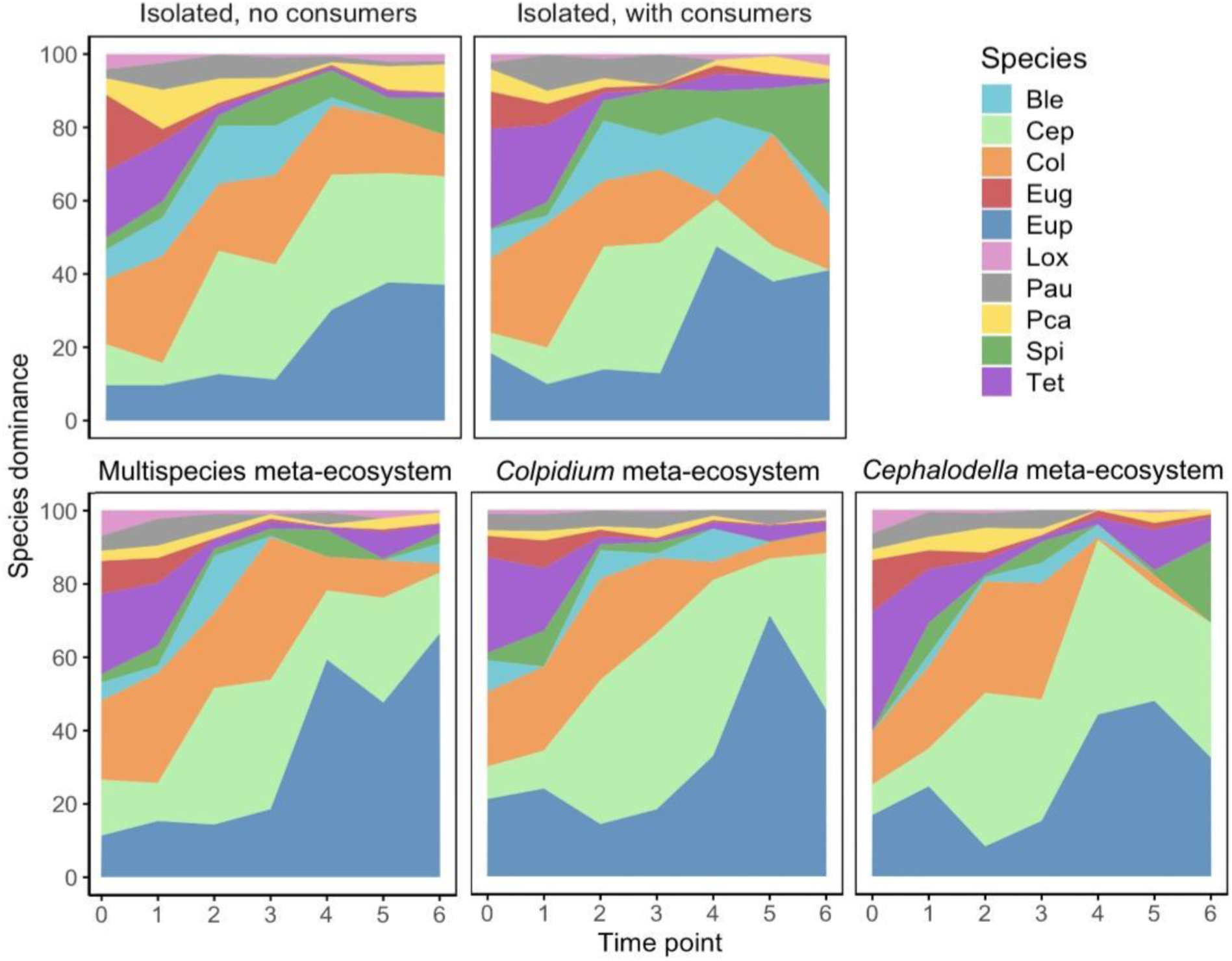
Time series of species dominance in the multispecies community in ecosystem 1 across isolated controls (top row) and meta-ecosystem treatments with consumer mobility (bottom row). Dominance is based on species biomass. Variation across replicate treatments is omitted for clarity (*n* = 5), but the same data with variability measures included can be observed in the Supplementary Material (Fig. S2). Ble = *Blepharisma* sp.; Cep = *Cephalodella* sp.; Col = *Colpidium* sp.; Eug = *Euglena gracilis*; Eup = *Euplotes aediculatus*; Lox = *Loxocephalus* sp.; Pau = *Paramecium aurelia*; Spi = *Spirostomum teres*; Tet = *Tetrahymena* cf. *pyriformis*.

### Consumer mobility and composition of the connected ecosystem affect consumer growth and reproductive output

The mobility of consumers and the composition of the connected ecosystem resulted in significant differences in the growth of the *Daphnia* consumers across the experiment (Fig. 4). *Daphnia* in the multispecies meta-ecosystem (pink lines) grew significantly larger than the isolated control with *Daphnia* (Fig. 4; grey lines; t = −3.381, df = 15, p = 0.019) and the *Colpidium* meta-ecosystems (blue lines; t = −6.624, df = 15, p < 0.001). While *Daphnia* in the multispecies meta-ecosystem also tended to be larger than the *Cephalodella* meta-ecosystem, the difference was not significant (yellow lines: t = 2.759, df = 15, p = 0.063). The size of *Daphnia* in the *Cephalodella* meta-ecosystem was not significantly different from the isolated control with *Daphnia* (t = −0.660, df = 15, p = 0.910) but was significantly larger than *Daphnia* in the *Colpidium* meta-ecosystem (t = −4.100, df = 15, p = 0.005).

**Figure 4.**
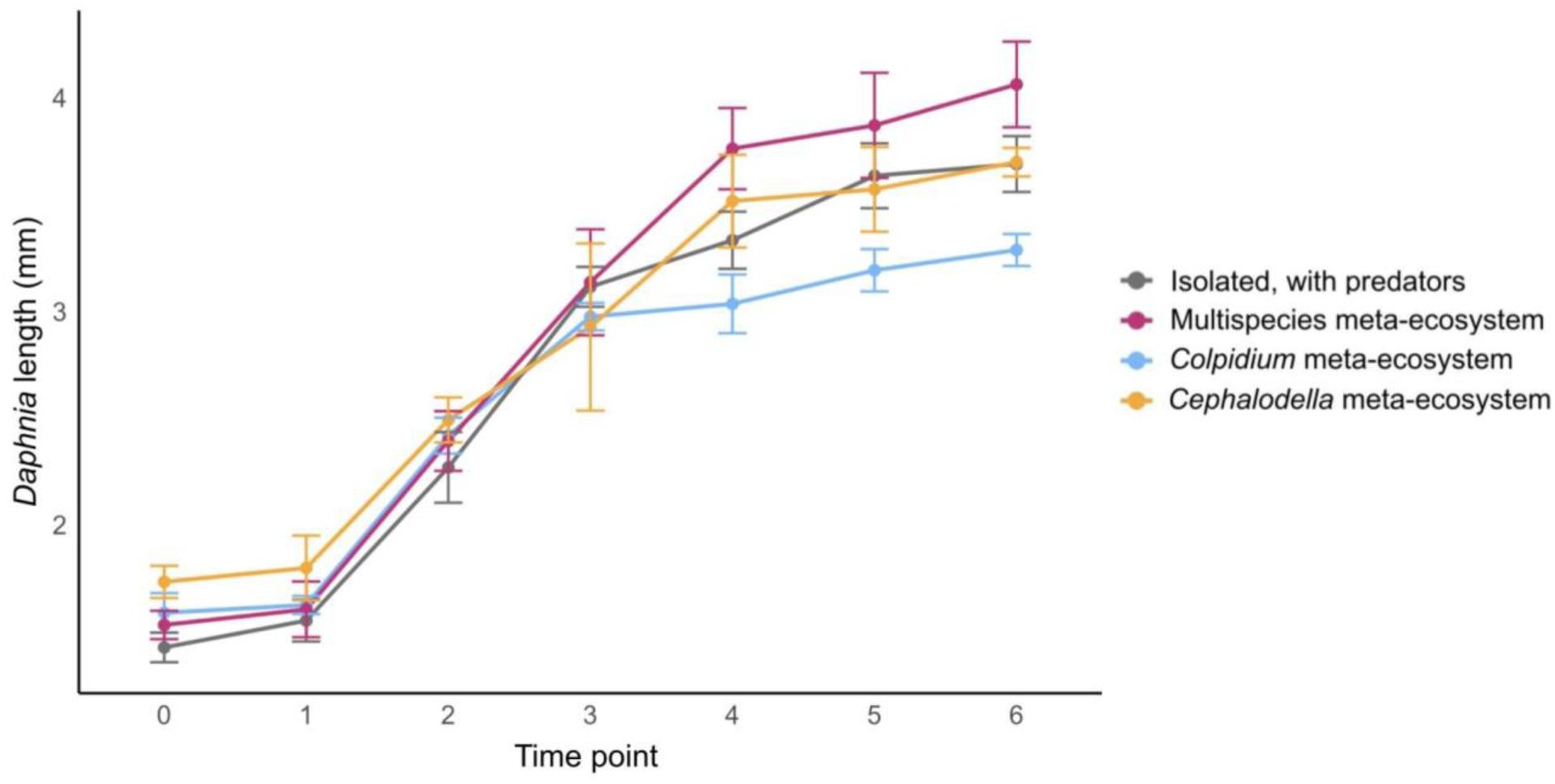
*Daphnia* growth across the experiment expressed as changes in length (mm). Growth is shown for isolated ecosystems, where *Daphnia* are restricted to a single ecosystem (grey), versus when *Daphnia* are mobile in multispecies (pink), *Colpidium* (blue) and *Cephalodella* (yellow) meta-ecosystems. Points give the mean across replicates and bars give the 95% confidence intervals.

Similarly, we found the mobility of consumers and the composition of the connected ecosystem resulted in significant differences in the reproductive output of the *Daphnia* consumers (Fig. 5; F = 10.09, df = 3, p = < 0.001). Relative to the isolated ecosystem with consumers (grey boxplot), the reproductive output of *Daphnia* was significantly higher in the multispecies meta-ecosystem (pink boxplot; p = 0.047) and the *Cephalodella* meta-ecosystem (yellow boxplot; p < 0.001), but was not statistically different in the *Colpidium* meta-ecosystem (blue boxplot; p = 0.74). The reproductive output of *Daphnia* in the *Cephalodella* meta-ecosystem did not significantly differ from the multispecies meta-ecosystem (p = 0.28), but was significantly higher than the *Colpidium* meta-ecosystem (p = 0.005). There was no significant difference in output between the multispecies meta-ecosystem and the *Colpidium* meta-ecosystem (p = 0.25).

**Figure 5.**
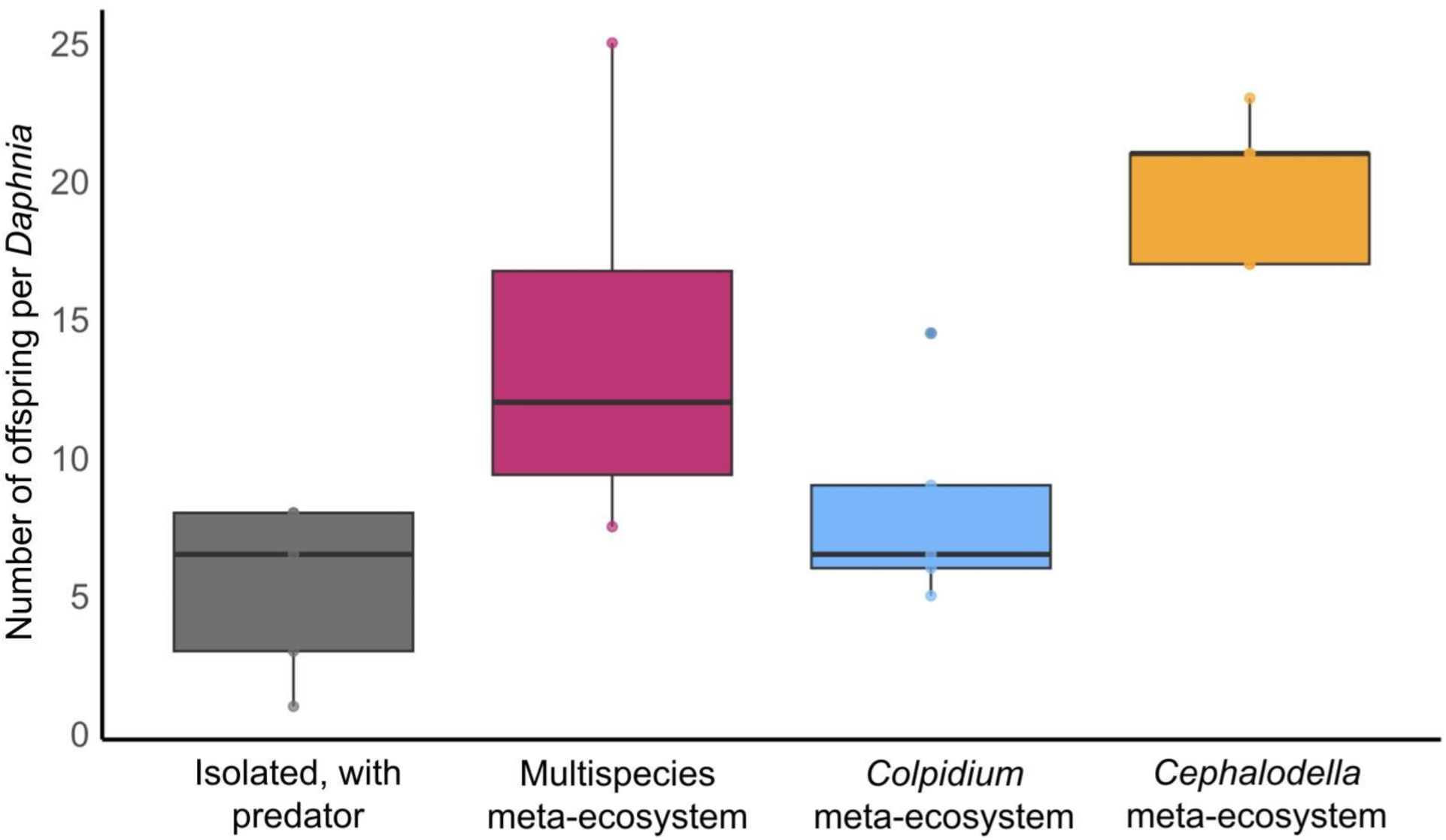
Average number of offspring produced per *Daphnia* when restricted to an isolated ecosystem (grey) versus when *Daphnia* are mobile in multispecies (pink), *Colpidium* (blue) and *Cephalodella* (yellow) meta-ecosystems. Boxplots show the first and third quartiles (box limits), median (midline), and 1.5× interquartile ranges (whiskers). Dots indicate the individual replicates.

## Discussion

We used a protist meta-ecosystem experiment to test how consumer mobility and the composition of the communities that consumers couple in space, influence community dynamics (species dominance), ecosystem functions (biomass) and affect the growth and reproductive output of consumers themselves. Our results demonstrate that consumer mobility can significantly reduce the negative impact of consumers on ecosystem biomass, and positively impact the growth and reproductive output of consumers. Notably, however, we found community composition of the connected ecosystem (ecosystem 2) could significantly mediate the direction and magnitude of these effects. Finally, we found consumer mobility consistently altered species dominance rankings, regardless of community composition in the connected ecosystem, by enabling the persistence of a local competitor of the mobile species. Ultimately, our study provides novel experimental evidence that consumer mobility is a profound driver of ecological dynamics, with its effects shaped by the composition of the ecosystems it couples. As global change alters species movement patterns and reshapes local communities, our results suggest the potential for cascading ecological effects across spatially connected ecosystems.

### Consumer mobility and composition of the connected ecosystem influence ecosystem biomass

Our results provide experimental evidence that consumer mobility can significantly influence ecosystem biomass. When consumers moved between two ecosystems with the same community composition, total biomass was significantly higher in the focal ecosystem relative to the immobile consumer control, where consumers were restricted to a single ecosystem. This baseline result suggests that consumer mobility, by allowing access to multiple ecosystems and thus, alleviating constant top-down pressure in any single ecosystem, can reduce the intensity of negative consumer impacts on prey biomass. This finding aligns with evidence from the field and supports theoretical predictions that consumer mobility can reduce the intensity of consumptive impacts on prey populations (Holt 1987, McCann et al., 2005, Peller et al. 2022)— an idea that, despite its theoretical basis, has been lacking robust experimental validation. It also aligns with studies identifying dispersal syndromes, whereby the ecosystem level effects (decomposition) of dispersing individuals was up to three times higher than of non-dispersing individuals in coupled two-patch systems (Little et al. 2019). Here, we show that such ecosystem effects are not only a behavioral consequence, but are shaped and affected by community composition. Our study demonstrates the critical role of consumer mobility in shaping ecosystem biomass and offers a versatile experimental approach that complements theoretical perspectives, enabling deeper exploration of these ecological processes.

Critically, we observed community composition of the connected ecosystem played a significant role in mediating the effects of mobile consumers on the focal ecosystem. Specifically, the effects of mobile consumers connected to a *Cephalodella* ecosystem (larger, more difficult prey to consume) did not significantly differ from the effects of immobile consumers restricted to the isolated control. In contrast, the effects of mobile consumers connected to a *Colpidium* ecosystem (smaller, less difficult prey to consume) tended to further weaken consumer effects relative to the multispecies meta-ecosystem, albeit not significantly. These findings supported our expectation that being connected to an ecosystem with relatively difficult consumption opportunities would result in strong consumptive effects in the focal ecosystem, likely because consumers were relatively starved following their visits to the *Cephalodella* ecosystem. Indeed, studies have shown organisms can have significantly higher consumption rates following periods of starvation versus satiation (Cronin and Hay 1996, Xuwang et al. 2011) or alter other behaviours to promote food acquisition (Vadas et al. 1994). Our findings build on this previous work by suggesting unfavorable feeding conditions in one ecosystem can amplify consumptive effects in a more favorable feeding ecosystem, leading to effects comparable to continuous consumer presence. Ultimately, our study highlights how community composition can trigger spatially cascading ecological effects by influencing the feeding dynamics of mobile organisms.

### Consumer mobility mediates local species dominance

Our study provides experimental evidence that consumer mobility can shape species dominance within the ecosystems they connect. In particular, we found that *Cephalodella sp.*, which could not persist long-term in the isolated control with *Daphnia*, was maintained as the second most dominant species across all meta-ecosystem treatments where consumers were mobile. Notably, the literature has long shown that *Daphnia* can both feed on ciliates and rotifers, and compete with rotifers for protist prey (Fenchel 1980, Bogdan and Gilbert 1987), with *Cephalodella* being competitively inferior to *Daphnia* (Gilbert 1985). It is, therefore, unsurprising that the constant presence of *Daphnia* in the isolated control resulted in the disappearance of *Cephalodella*, likely due to insufficient prey availability as *Daphnia* monopolized shared resources. Our findings suggest that consumer mobility, by periodically alleviating predation pressure and competition, can mediate competitive dynamics and facilitate the persistence of otherwise inferior competitors. These results support theoretical predictions that mobility can enhance coexistence by buffering species from competitive exclusion (Chesson 2000, Vincent et al. 1996, Peller et al. 2022). Overall, our findings highlight that consumer mobility can influence community composition and species dominance by creating opportunities for coexistence that would otherwise be unavailable in isolated systems.

### Consumer mobility and composition of the connected ecosystem influence consumer fitness

Our experiment demonstrated that consumer mobility, and community composition of the connected ecosystems, can significantly influence consumer growth and reproductive output. Specifically, *Daphnia* in the multispecies meta-ecosystem had significantly greater growth and produced significantly more offspring than *Daphnia* in the isolated control. This finding supports theoretical predictions that consumer mobility generally increases the overall biomass of the mobile consumers by enabling greater total consumption (e.g., McCann et al. 2005, Peller et al. 2022). Moreover, this finding aligns with field evidence, for instance, showing that fish that move between patches tracking abundant resources have significantly higher growth rates than those that remain within a single patch (Ruff et al. 2011), just as resource subsidies can infer greater growth and reproductive outputs by increasing resource availability across time (Tylianakis et al. 2004, Wright et al. 2013). In our experiment, moving between two ecosystems with similar community composition alleviated the negative impact of consumption on ecosystem biomass, and thus, consumers experienced greater prey availability across the meta-ecosystem conferring such growth and reproductive advantages. Importantly, these findings suggest that increased biomass resulting from mobile consumers coupling ecosystems can occur via both increased consumer growth rates and reproductive output.

Consistent with the biomass effects of consumer mobility, community composition in the connected ecosystem influenced consumer growth and reproductive output. In *Cephalodella* meta-ecosystems, growth of *Daphnia* did not significantly differ from the isolated control (echoing the biomass result), but reproductive output was significantly higher. Due to the strong negative impact of *Daphnia* on biomass in the focal ecosystem in *Cephalodella* meta-ecosystems, we expect *Daphnia* growth did not benefit from increased resource availability as it did in the multispecies meta-ecosystem. The less intuitive finding is that *Daphnia* in *Cephalodella* meta-ecosystems produced significantly more offspring than they did in the isolated control and the other meta-ecosystem treatments. One possible explanation, based on previous experimental findings, is that *Daphnia* which feed on lower quality prey items can invest in a greater number of eggs of smaller size (Frost et al. 2010, Bednarska 2022). As there is evidence to suggest rotifers are not a high-quality prey for *Daphnia* (Wickham et al. 1993), it is possible *Daphnia* in *Cephalodella* meta-ecosystems produced more, yet smaller offspring. Another possible explanation is that the visits to the *Cephalodella* ecosystem were short enough such that *Daphnia* were able to recover upon their return to the multispecies ecosystem, while investing more in reproduction than growth. Indeed, *Daphnia* have been observed to use energy reserves in the body to maintain reproduction when conditions are less favorable (Tessier et al. 1983), and to more efficiently use high quality foods when more favorable conditions return, potentially allowing quick recovery (Xuwang et al. 2011).

In *Colpidium* meta-ecosystems, *Daphnia* growth was significantly lower than the isolated control and the other meta-ecosystem treatments, and *Daphnia* reproductive output was not significantly different from the isolated control. Previous experiments have demonstrated a strong predation effect of *Daphnia* on *Colpidium* (Gounand et al. 2017), suggesting that *Daphnia* readily consume *Colpidium*. Our findings, however, suggest *Colpidium* may be a relatively low-quality prey source for *Daphnia*, negatively influencing their fitness relative to more diverse prey compositions. As growth and reproduction of Daphnia in *Colpidium* meta-ecosystems were reduced relative to *Cephalodella* meta-ecosystems, this suggests that exposure to a low-quality, easily consumed prey can have negative fitness consequences in meta-ecosystems compared to exposure to more difficult-to-consume prey. That is, by filling up on low-quality prey, consumers may have left the *Colpidium* ecosystem more satiated than when leaving the *Cephalodella* ecosystem, preventing increased feeding effort in the focal ecosystem, which is consistent with our biomass results (i.e., higher biomass in the focal ecosystems of *Colpidium* versus *Cephalodella* meta-ecosystems). Studies of subsidized ecosystems also support this expectation, as shown, for instance, by the negative effects of readily available and consumable fisheries waste on Cape gannet chicks (Fremillet et al. 2008).

### Implications and Conclusions

Our study represents an important step forward in experimental ecology, presenting a versatile meta-ecosystem set-up for testing how consumer mobility and community composition influence ecological dynamics. Despite highly controlled, spatially structured experiments being widely applied to test the effects of dispersal on ecosystems (Shurin 2001, Fronhofer et al. 2015, Grainger and Gilbert 2016, Gilarranz et al. 2017), experiments testing such predictions on non-dispersal movements are scarce. By incorporating community composition into our framework, we demonstrate how ecosystem diversity plays a pivotal role in shaping the effects of mobile consumers. These insights extend beyond theoretical applications, providing valuable perspectives for conservation and management efforts, where maintaining connectivity and considering the diversity of interconnected ecosystems could promote ecosystem functions and enhance biodiversity. Altogether, our study underscores the importance of fostering greater interaction between theoretical and experimental approaches to deepen our understanding of consumer mobility. By advancing our understanding of how consumer mobility interacts with community composition to shape ecological outcomes, our work provides a foundation for future research exploring the dynamic interplay between movement, biodiversity, and ecosystem function in a rapidly changing world.

## Acknowledgement

Funding is from the Swiss National Science Foundation (Grant 310030_197410).

## Literature Cited

Abbas, F., Merlet, J., Morellet, N., Cwehwyswn, H., Hewison, A.J., Cargnelutti, B., et al. (2012). Roe deer markedly alter forest nitrogen and phosphorous budgets across Europe. Oikos, 121: 1271–1278.

Altermatt, F., Fronhofer, E.A., Garnier, A., Giometto, A., Hammes, F., Klecka, J., et al. (2015). Big answers from small worlds: A user’s guide for protist microcosms as a model system in ecology and evolution. Methods in Ecology and Evolution, 6: 218–231.

Bates, D., Mächler, M., Bolker, B.M., and Walker, S.C. (2015). Fitting linear mixed-effects models using lme4. Journal of Statistical Software, 67: 1–48.

Bandara, K., Varpe, O, Wijewardene, L., Tverberg, V., and Eiane, K. (2021). Two hundred years of zooplankton vertical migration research. Biological Reviews, 96: 1547–1589.

Bauer, S. and Hoye, B.J. (2014). Migratory animals couple biodiversity and ecosystem functioning worldwide. Science, 344: 1242552.

Bednarska, A. (2022). Food quantity and quality shapes reproductive strategies of *Daphnia*. Ecology and Evolution, 12: e9163.

Bogdan, K.G. and Gilbert, J.J. (1987). Quantitative comparison of food niches in some freshwater zooplankton. Oecologia, 72: 331–340.

Chesson, P. (2000). Mechanisms of maintenance of species diversity. Annual Review of Ecology Evolution and Systematics, 31: 343–366.

Cortes, C., Vapnik, V., and Saitta, L. (1995). Support vector networks. Machine Learning, 20: 273–297.

Cronin, G. and Hay, M.E. (1996). Susceptibility to herbivores depends on recent history of both the plant and animal. Ecology, 77: 1531–1543.

Deacy, W.W., Armstrong, J.B., Leacock, W.B., Robbins, C.T., Gustine, D.D., Ward, E.J., et al. (2017). Phenological synchronization disrupts trophic interactions between Kodiak brown bears and salmon. PNAS, 114: 10432–10437.

DeBiase, A.E., Sanders, R.W., and Porter, K.G. (1990). Relative nutritional value of ciliate protozoa and algae as food for *Daphnia*. Microbial Ecology, 19: 199–210.

Dimitriadou, E., Hornik, K., Leisch, F., Meyer, D., and Maintainer, A. W. (2006). Misc Functions of the Department of Statistics (e1071), TU Wien.

Donadi, S., Austin, Å.N., Bergström, U., Eriksson, B.K., Hansen, J.P., Jacobson, P., et al. (2017). A cross-scale trophic cascade from large predatory fish to algae in coastal ecosystems. Proceedings of the Royal Society B, 284: 20170045.

Duckworth, J., Jager, T., and Ashauer, R. (2019). Automated, high-throughput measurement of size and growth curves of small organisms in well plates. Scientific Reports, 9: 10.

Fenchel, T. (1980). Suspension feeding in ciliated protozoa: functional response and particle size selection. Microbial Ecology, 6: 1–11.

Fremillet, D., Pichegru, L., Kuntz, G., Woakes, A.G., Wilkinson, S., Crawford, R.J., et al. (2008). A junk-food hypothesis for gannets feeding on fishery waste. Proceedings of the Royal Society B, 275: 1149–1156.

Fox, J. and Weisberg, S. (2011). An {R} companion to applied regression. – Sage.

Fronhofer, E.A., Klecka, J., Melián, C.J., and Altermatt, F. (2015). Condition-dependent movement and dispersal in experimental metacommunities. Ecology Letters, 18: 954–963.

Frost, P.C., Ebert, D., Larson, J.H., Marcus, M.A., Wagner, N.D., and Zalewski, A. (2010). Transgenerational effects of poor elemental food quality on *Daphnia magna*. Oecologia, 162, 865–872.

Fussman, G.F., Ellner, S.P., Shertzer, K.W., and Hairston Jr, N.G. (2000). Crossing the hopf bifurcation in a live predator-prey system. Science, 290: 1358–1360.

Giacomuzzo, E., Peller, T., Gounand, I., and Altermatt, F. (2024). Ecosystem size mediates the effects of resource flows on species diversity and ecosystem functions at different scales. Ecology and Evolution, 14: e70709.

Gilarranz, L.J., Rayfield, B., Liñan-Cembrano, G., Bascompte, J., and Gonzalex, A. (2017). Effects of network modularity on the spread of perturbation impact in experimental metapopulations. Science, 357: 199–201.

Gilbert, J.T. (1985). Competition between Rotifers and Daphnia. Ecology, 66: 1943–1950.

Giometto, A., Altermatt, F., Carrara, F., and Rinaldo, A. (2013). Scaling body size fluctuations. PNAS, 110: 4646–4650.

Gounand, I., Harvey, E., Ganesanandamoorthy, P., and Altermatt, F. (2017). Subsidies mediate interactions between communities across space. Oikos, 126: 972–979.

Gounand, I., Little, C.J., Harvey, E., and Altermatt, F. (2018a). Cross-ecosystem carbon flows connecting ecosystems worldwide. Nature Communications, 9: 4825.

Gounand, I., Harvey, E., Little, C.J. and Altermatt, F. (2018b). Meta-ecosystems 2.0: rooting the theory into the field. Trends in Ecology & Evolution, 33: 36–46.

Grainger, T.N. and Gilbert, B. (2016). Dispersal and diversity in experimental metacommunities: linking theory and practice. Oikos, 125: 1213–1223.

Green, A.J., Lovas-Kiss, A., Reynolds, C., Sebastian-Gonzalez, E., Silva, G.G., van Leeuwen, C.H.A., and Wilkinson, D.M. (2023). Dispersal of aquatic and terrestrial organisms by waterbirds: a review of current knowledge and future priorities. Freshwater Biology, 68: 173–190.

Guzman, L.M., Germain, R.M., Forbes, C., Straus, S., O’Connor, M.I., Gravel, D. et al. (2019). Towards a multi-trophic extension of metacommunity ecology. Ecology Letters, 22: 19–33.

Holt, R.D. (1987). Prey communities in patchy environments. Oikos, 50: 276–290.

Jack, J. D. and Gilbert, J.J. (1993). Susceptibilities of different-sized ciliates to direct suppression by small and large cladocerans. Freshwater Biology, 29: 19–29.

Kauffman, M.J., Aikens, E.O., Esmaeili, S., Kaczensky, P., Middelton, A., Monteith, K.L., et al. (2021). Causes, consequences, and conservation of ungulate migration. Annual Review of Ecology, Evolution, and Systematics, 52: 453–478.

Kremen, C., Williams, N.M., Aizen, M.A., Gemmill-Herren, B., LeBuhn, G., Minckley, R., et al. (2007). Pollination and other ecosystem services produced by mobile organisms: a conceptual framework for the effects of land-use change. Ecology Letters, 10: 299–314.

Lawler, S.P. (1998). Ecology in a Bottle Using Microcosms to Test Theory. In Experimental Ecology: Issues and Perspectives. Oxford University Press: Oxford.

Leibold, M.A. (1989). Resource edibility and the effects of predators and productivity on the outcome of trophic interactions. The American Naturalist, 134: 922–949.

Lenth, R.V. (2022). Emmeans: Estimated Marginal Means, aka Least-Squares Means.

Leroux, S.J. and Schmitz, O.J. (2025). Integrating network and meta-ecosystem models for developing a zoogeochemical theory. Ecology Letters, 28: e70076.

Little, C.J., Fronhofer, E.A., and Altermatt, F. (2019). Dispersal syndromes can impact ecosystem functioning in spatially structured freshwater populations. Biology Letters, 15: 20180865.

Lundberg, J. and Moberg, F. (2003). Mobile link organisms and ecosystem functioning: implications for ecosystem resilience and management. Ecosystems, 6: 0087–0098.

Massol, F., Gravel, D., Mouquet, N., Cadotte, M.W., Fukami, T., and Leibold, M.A. (2011). Linking community and ecosystem dynamics through spatial ecology. Ecology Letters, 14: 313–323.

McCann, K.S., Rasmussen, J.B. and Umbanhowar, J. (2005). The dynamics of spatially coupled food webs. Ecology Letters, 8: 513–523.

McMahon, J.W. and Rigler, F.H. (1965). Feeding rate of *Daphnia magna* straus in different foods labeled with radioactive phosphorus. Limnology and Oceanography, 10: 105–113.

Meyer, J.L., Schultz, E.T., and Helfman, G.S. (1983). Fish schools: an asset to corals. Science, 220: 1047–1049.

Oksanen, J., Simpson, G., Blanchet, F., Kindt, R., Legendre, P., Minchin, P., et al. (2024). Vegan: Community Ecology Package. R package version 2.6-8, <https://CRAN.R-project.org/package=vegan>.

Pace, M.L. and Funke, E. (1991). Regulation of planktonic microbial communities by nutrients and herbivores. Ecology, 72: 904–914.

Peller, T., Andrews, S., Leroux, S.J. and Guichard, F. (2021). From marine metacommunities to meta-ecosystems: examining the nature, scale, and significance of resource flows in benthic marine environments. Ecosystems, 24: 1239–1252.

Peller, T., Marleau, J.N. and Guichard, F. (2022). Traits affecting nutrient recycling by mobile consumers can explain coexistence and spatially heterogeneous trophic regulation across a meta-ecosystem. Ecology Letters, 25: 440–452.

Peller, T., Guichard, F., and Altermatt, F. (2023). The significance of partial migration for food web and ecosystem dynamics. Ecology Letters, 26: 3–22.

Peller, T., Gounand, I., and Altermatt, F. (2024). Resource flow network structure drives meta-ecosystem function. The American Naturalist, 204: 546–560.

Pennekamp, F., Schtickzelle, N., and Petchey, O.L. (2015). BEMOVI, software for extracting behaviour and morphology from videos, illustrated with analyses of microbes. Ecology and Evolution, 5: 2584–2595.

Pennekamp, F., Pontarp, M., Tabi, A., Altermatt, F., Alther, R., Choffat, Y., et al. (2018). Biodiversity increases and decreases ecosystem stability. Nature 563: 109–112.

Petchey, O.L. (2001). Prey diversity, prey composition, and predatory population dynamics in experimental microcosms. Journal of Animal Ecology, 69: 874–882.

Polis, G.A., Anderson, W.B. and Holt, R.D. (1997). Toward an integration of landscape and food web ecology: the dynamics of spatially subsidized food webs. Annual Review of Ecology, Evolution, and Systematics, 28: 289–316.

Post, D.M., Conners, M.E., and Goldberg, D.S. (2000). Prey preference by a top predator and the stability of linked food chains. Ecology, 81: 8–14.

Rooney, N., McCann, K.S., and Moore, J.C. (2008). A landscape theory for food web architecture. Ecology Letters, 11: 867–881.

Rosen, D.A.S. and Trites, A.W. (2000). Pollock and the decline of Stellar sea lions: testing the junk-food hypothesis. Canadian Journal of Zoology, 78: 1243–1250.

Rosenberg, K.V., Dokter, A.M., Blancher, P.J., Sauer, J.R., Smith, A.C., Smith, P.A., et al. (2019). Decline of the North American avifauna. Science, 366: 120–124.

de Roos, A.M., McCauley, E., and Wilson, W.G. (1998). Pattern formation and the spatial scale of interaction between predators and prey. Theoretical Population Biology, 53: 108–130.

Ruff, C.P., Schindler, D.E., Armstrong, J.B., Bentley, K.T., Brooks, G.T., Holtgrieve, G.W., et al. (2011). Temperature-associated population diversity in salmon confers benefits to mobile consumers. Ecology, 92: 2073–2084.

Schneider, C.A., Rasband, W.S., and Eliceiri, K.W. (2012). NIH Image to ImageJ: 25 years of image analysis. Nature Methods, 9: 670–675.

Shurin, J.B. (2001). Interactive effects of predation and dispersal on zooplankton communities. Ecology, 82: 3404–3416.

Srivastava, D.S., Kolasa, J., Bengtsson, J., Gonzalez, A., Lawler, S.P., Miller, T.E., et al. (2004). Are natural microcosms useful model systems for ecology? Trends in Ecology and Evolution, 19: 379–384.

Tessier, A.J., Henry, L.L., Goulden, C.E., and Durand, M.W. (1983). Starvation in daphnia: energy reserves and reproductive allocation. Limnology and Oceanography, 28: 667–676.

Tylianakis, J.M., Didham, R.K., and Wratten, S.D. (2004). Improved fitness of aphid parasitoids receiving resource subsidies. Ecology, 85: 658–666.

Vadas, R.L., Burrows, M.T., and Hughes, R.N. (1994). Foraging strategies of dogwhelks, *Nucella lapillus* (L): interacting effects of age, diet, and chemical cues to threat of predation. Oecologia, 100: 439–450.

Vincent, T.L.S., Scheel, D., Brown, J.S., and Vincent, T.L. (1996). Trade-offs and coexistence in consumer-resource models: it all depends on what and where you eat. American Naturalist, 148: 1038–1058.

Wickham, S.A. and Gilbert, J.J. (1991). Relative vulnerabilities of natural rotifer and ciliate communities to cladocerans: laboratory and field experiments. Freshwater Biology, 26: 77–86.

Wickham, S.A., Gilbert, J.J., and Berninger, U.-G. (1993). Effects of rotifers and ciliates on the growth and survival of *Daphnia*. Journal of Plankton Research, 15: 317–334.

Williams, S.H., Steenweg, R., Hegel, T., Russel, M., Dervieux, D., and Hebblewhite, M. (2021). Habitat loss on seasonal migratory range imperils an endangered ungulate. Ecological Solutions and Evidence, 2: e12039.

Wright, A.N., Piovia-Scott, J., Spiller, D.A., Takimoto, G., Yang, L.H., and Schoener, T.W. (2013). Pulses of marine subsidies amplify reproductive potential of lizards by increasing individual growth rate. Oikos, 122: 1496–1504.

Xuwang, Y., Shasha, Z., Jian, H., and Pengfei, L. (2011). Effects of food quality and starvation on the optimal foraging behaviour of *Daphnia magna* (Cladocera). Acta Ecologica Sinica, 31: 328–333.

Zöllner, E., Santer, B., Boersma, M., Hoppe, H.-G., and Jürgens, K. (2003). Cascading predation effects of *Daphnia* and copepods on microbial food web components. Freshwater Biology, 48: 2174–2193.

