## Supplementary Information for "Food-web coupling by mobile consumers has individual to ecosystem level effects"

**Total biomass in ecosystem 2 across the experiment**

To assess whether the effects in ecosystem 1 could be driven by resource availability in ecosystem 2, rather than community composition, we tested whether biomass in ecosystem 2 significantly different between multispecies, *Colpidium*, and *Cephalodella* meta-ecosystems. Specifically, we ran a linear mixed effects model testing the interactive influence of meta-ecosystem type and continuous time on biomass in ecosystem 2, with replicate as a random effect. We performed post hoc analyses to evaluate the significance of pairwise differences between treatments (with Tukey adjusted p-values).

The model revealed no significant differences in biomass in ecosystem 2 among different meta-ecosystem types (Fig. S1). Post hoc pairwise comparisons showed no significant differences in biomass between the multispecies and *Colpidium* meta-ecosystems ($t = -0.815$, $df = 12$, $p = 0.701$), between the multispecies and *Cephalodella* meta-ecosystems ($t = 1.160$, $df = 12$, $p = 0.498$), or between the *Colpidium* and *Cephalodella* meta-ecosystems ($t = 1.976$, $df = 12$, $p = 0.161$). These results suggest that resource availability in ecosystem 2 did not differ significantly among meta-ecosystem types, reinforcing the interpretation that observed effects in ecosystem 1 were driven primarily by community composition rather than differences in resource supply.

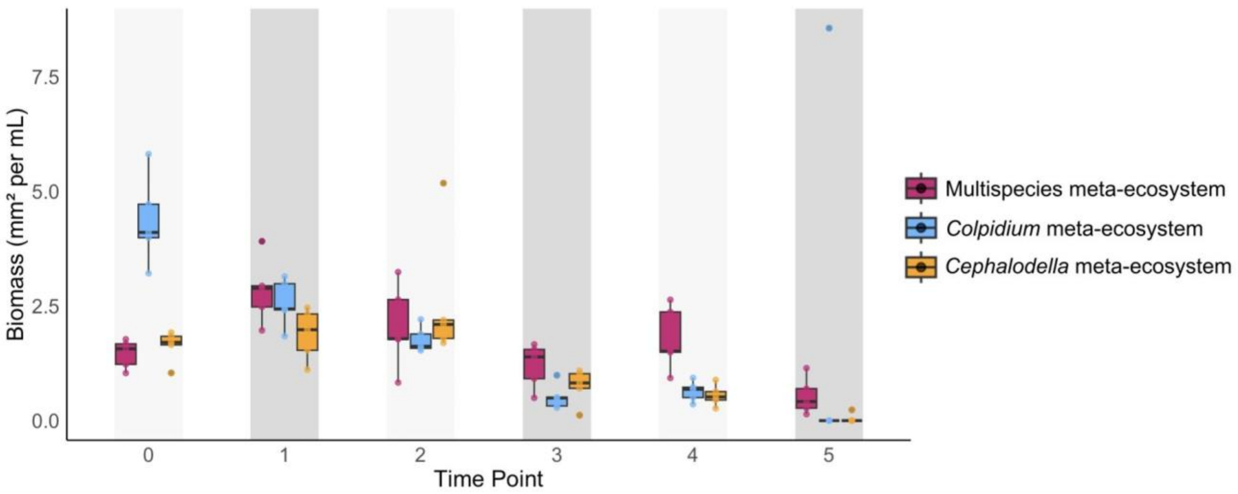


**Figure S1. Total biomass in ecosystem 2 across the experiment.** Ecosystem 2 in the multispecies meta-ecosystem (pink) consists of the same mixed multispecies community as all treatments have in ecosystem 1. Ecosystem 2 in the *Colpidium* meta-ecosystem (blue) consists of a *Colpidium* sp. monoculture. Ecosystem 2 in the *Cephalodella* metaecosystem (yellow) consists of a *Cephalodella* sp. monoculture. Time points with light grey shading show the biomass in ecosystem 2 directly before *Daphnia* consumers were transferred to ecosystem 2. Whereas time points with dark grey shading show the biomass in ecosystem 2 directly after *Daphnia* consumers were removed from ecosystem 2 and transferred to ecosystem 1.

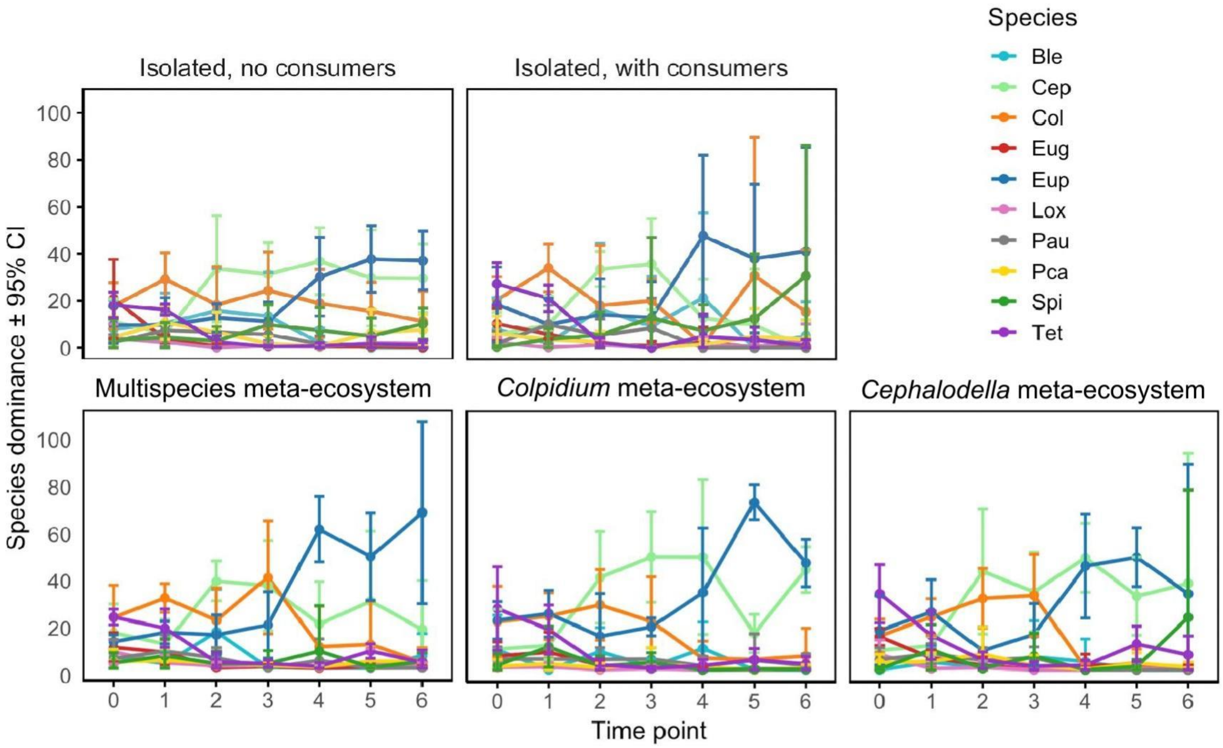


**Figure S2.** Time series of species dominance in the multispecies community in ecosystem 1 across isolated controls (top row) and meta-ecosystem treatments with consumer mobility (bottom row). Dominance is based on species biomass, and the plot shows the mean values with 95% confidence intervals to represent variation across replicate treatments. Ble = *Blepharisma sp.*; Cep = *Cephalodella sp.*; Col = *Colpidium* sp.; Eug = *Euglena gracilis*; Eup = *Euplotes aediculatus*; Lox = *Loxocephalus* sp.; Pau = *Paramecium aurelia*; Spi = *Spirostomum teres*; Tet = *Tetrahymena* cf. *pyriformis*.

**Table S1.** Effect of community composition in ecosystem 2 on log response ratio of biomass in ecosystem 1 across time points. Analysis of deviance (type II tests) on the generalized linear models. Gaussian distributions were used for models. The table reports the likelihood ratio chi-squared (LR Chisq), degrees of freedom (DF), and the respective p-values for the significance tests.

| | **LR Chisq** | **DF** | **p-value** |
|---|---|---|---|
| Community composition | 20.574 | 3 | **0.000129** |
| Time | 23.650 | 1 | **<0.0001** |
| Community composition x time | 10.930 | 3 | **0.012112** |
